# “Altered CD8^+^ T cell associated aging gene signature in the peripheral blood of patients with Alzheimer’s disease”

**DOI:** 10.1101/2022.09.29.510171

**Authors:** Juan J. Young, Hong-Jai Park, Minhyung Kim, Jennefer Par-Young, Hugh H. Bartlett, Hye Sun Kim, Min Sun Shin, Serhan Unlu, Richard Bucala, Christopher H. Van Dyck, Heather G. Allore, Adam P. Mecca, Sungyong You, Insoo Kang

## Abstract

**INTRODUCTION:** Effector memory (EM) CD8^+^ T cells have been associated with poor cognition in Alzheimer’s disease (AD). Our lab recently discovered an age-associated gene expression signature of IL-7 receptor alpha (IL-7Rα)^low^ EM CD8^+^ T cells. We hypothesized that individuals with AD have altered levels of this IL-7Rα^low^ aging gene expression.

**METHODS:** Forty genes associated with IL-7Rα^low^ EM CD8^+^ T cells, AD, or memory, were analyzed in peripheral blood of participants with normal cognition, mild cognitive impairment, and dementia by qPCR.

**RESULTS:** Of the eight genes that were found to be differentially expressed based on clinical diagnosis, 5 genes (62.5%) were IL-7Rα^low^ aging genes. Principal component analysis revealed 3 clusters of participants with dementia which had distinct expression levels of IL-7Rα^low^ aging genes and cognitive function.

**DISCUSSION:** Our findings support the possible relationship of the IL-7Rα^low^ EM CD8^+^ T cell aging signature with cognition in individuals with dementia due to AD.

## 1. INTRODUCTION

Alzheimer’s disease (AD) is a chronic progressive neurodegenerative disorder and the most common cause of dementia, accounting for approximately 60-80% of all dementia cases [1]. Amyloid-β (Aβ), a peptide derived from the amyloid precursor protein (APP) and known to assemble into extracellular amyloid plaques, is hypothesized to initiate AD pathogenesis by stimulating the formation of neurofibrillary tangles and contributing to the development of a cytotoxic environment prone to neurodegeneration [2]. Successive stressful events early in life including infections, ischemia, free radicals, and metabolic insults (*e*.*g*., overnutrition) are thought to result in a chronic, low-intensity inflammatory state that stimulates pathological Aβ accumulation and the development of the AD dementia syndrome [3, 4]. Previous studies have focused primarily on the innate immune system, including microglial cells which can be activated by Aβ, as a direct approach to evaluating neuroinflammation in the central nervous system (CNS) [5]. In contrast, few have investigated the potential role of adaptive immunity, including T cells, on AD pathology as part of a dysregulated systemic immune response resulting from harmful interactions between CNS immune cells and the peripheral immune system [6, 7].

Aging is the strongest risk factor for AD [1]. In human T cells, probably the most prominent change with aging is the expansion of effector memory (CCR7^-^, EM) CD8^+^ T cells, which include CD45RA^+^ and CD45RA^−^ populations (**note**: hereafter EM indicates both CD45RA^+^ and ^−^ populations unless specified), in peripheral blood [8, 9]. One potential contributor to AD pathology includes cytotoxic and senescent T cell populations that may interact with the CNS through disruptions in the blood brain barrier [7, 10]. Previously, our lab found an age-associated expansion of peripheral cytotoxic human effector memory (EM, CCR7^-^CD45RA^+/-^) CD8^+^ T cells expressing low levels of IL-7 receptor alpha chain (IL-7Rα or CD127)^low^, which have distinct characteristics including effector molecules, transcription factors, and DNA methylation profiles [10-13]. Such T cell expansion, which is likely driven in part by repetitive immune stimulation over a lifetime [14], contributes to age-associated transcriptomic changes in human peripheral blood cells. About 15% of the age□associated genes (231/1,497) reported by a meta□analysis study of human peripheral blood from approximately 15,000 individuals corresponded to differentially expressed genes in IL-7Rα^low^ EM CD8^+^ T cells [15].

A recent study reported the expansion of CD45RA^+^ EM CD8^+^ T cells (T_EMRA_) in the cerebrospinal fluid of patients with dementia or mild cognitive impairment (MCI) due to AD [16]. The clonal expansion of these cells in peripheral blood was found to also correlate with cognitive scores. Importantly, such cells are mostly IL-7Rα^low^ EM CD8^+^ T cells, implying the possible relationship of such cell expansion with transcriptomic alterations in the peripheral blood of individuals with AD. Thus, the present study was conducted to determine if an altered IL-7Rα^low^ EM CD8^+^ T cell associated aging gene signature occurs in peripheral blood of AD participants compared to cognitively normal (CN) persons.

## 2. METHODS

### 2.1. Participant Characteristics

A total of 121 whole blood samples from participants recruited by the Yale ADRC were requested and obtained for processing and transcriptomic analysis (Table 1). Participants included in this study were organized into the following clinical groups based on the consensus diagnosis: CN participants, MCI due to AD, and probable dementia due to AD. MCI and probable dementia due to AD are defined according to the National Institute on Aging – Alzheimer’s Association (NIA-AA) guidelines [17, 18]. Clinical data from each participant, including biomarkers and cognitive testing scores, were utilized to produce a consensus diagnosis based on review by a multidisciplinary panel of experts from the Yale ADRC. Severity and staging were assessed using clinical rating scales including the Montreal Cognitive Assessment (MoCA) [19], and Washington University’s Clinical Dementia Rating (CDR) scale [20]. Of these participants, 38 were CN, 40 were diagnosed with MCI due to AD, and 43 were diagnosed with dementia due to AD. Though the three clinical groups were age-matched and were similar in distribution in male and female participants, they were found to have an unequal distribution of participants based on the following racial categories: White, Black, American Indian/Native Alaskan, Asian, or Unknown (i.e., participants that declined to identify with any single race).

### 2.2. Selecting Genes of Interest

Forty genes of interest were chosen to be analyzed based on publicly available AD and memory transcriptomic datasets (GSE140829, and GSE63060 and GSE63061 for AD; GSE127711 for memory) as well as the Molecular signatures database (MSigDB) [21-24]. See Supplementary Methods 1.2 and Supplementary Table S1 for more details.

### 2.3. RNA Isolation and Complementary DNA synthesis

RNA was isolated from peripheral whole blood stored in heparinized tubes at −80°C using a modified QIAGEN RNeasy Kit protocol (see Supplementary Materials). RNA templates were then utilized for complimentary DNA (cDNA) synthesis utilizing the iScript cDNA synthesis kit protocol (Bio-Rad).

### 2.4. Quantitative Polymerase Chain Reaction

Target gene expression was measured by quantitative polymerase chain reaction (qPCR) analysis by producing 10 μl reaction mixtures containing cDNA, SYBR Green Supermix (Bio-Rad) and target gene primers at 1 μM concentrations (see Table S1. in supporting information for target gene primer RNA nucleotide sequences). The reaction mixture was initially denatured at 95°C for 3 minutes and then underwent 40 cycles of the following: denatured for 15 seconds at 95°C then annealing, extension, and read fluorescence for 45 seconds at 60°C using the CFX384 Touch Real-Time PCR Detection System (Bio-Rad). The expression levels for each gene were then calculated per the 2^-ΔΔC^T equation [25].

### 2.5. Data Processing

Approximately 2.6% of all expression fold change values comprising the current transcriptomic dataset were considered missing completely at random. Using Bioconductor’s “pcaMethods” software, missing values were imputed utilizing probabilistic principal component analysis (PPCA), a method of multiple imputation based on a probabilistic model created via a maximum likelihood estimation approach [26, 27]. ComBat, a program in the Bioconductor’s “sva” software suite that utilizes empirical Bayes regression to adjust for and correct batch effects [28, 29] was used to process the transcriptomic dataset generated from the RT-qPCR analysis.

### 2.6. Statistical Analyses

One-way ANOVA and Pearson’s chi-squared tests were performed as part of a descriptive analysis of ADRC participant demographic characteristics of age and sex, respectively. Due to the limited participation of non-White persons, analyses adjusting for race were conducted by labeling participants as either “non-Hispanic White” or “Other” with the latter group comprised of Hispanic and non-White participants. One-way ANOVA testing was completed to determine if there were differences in mean gene expression among the three clinical groups. General linear models (GLM) were also conducted to analyze differences in relative gene expression levels utilizing estimated least-square means (LSM) generated after adjusting for age, sex, and race. Fisher’s exact test was performed to calculate significance of overlap between 40 genes and genesets. Welch’s *t*-test was conducted to determine if there were differences in cognitive scores between dementia subgroup clusters. Data was processed and analyzed using IBM SPSS Statistics for Windows, Version 28.0, released in 2021, Armonk, NY: IBMCorp, R version 1.4.1106, and GraphPad Prism version 8.0.0 for Windows, GraphPad Software, San Diego, California USA.

## 3. RESULTS

### 3.1. A Systematic Selection of AD Associated Aging Genes

To select AD associated aging genes, we performed a systematic gene selection process on a large set of transcriptome data derived from the public domain geneset database and the published aging gene signature. In addition to the top 9 IL-7Rα^low^ EM CD8^+^ T cell aging genes from our previous report [15], we analyzed the rest of the 231 aging signature genes of IL-7Rα^low^ EM CD8^+^ T cells reported to be associated with or without AD in MsigDB and/or 3 peripheral blood transcriptomic studies of patients with AD (at least 2 of 3 analyses including GSE140829, GSE63060 and GSE63061). We also included 3 common genes between IL-7Rα^low^ EM CD8^+^ T cell associated aging genes and peripheral blood genes reported to be associated with short-term memory (GSE127711). We further included 18 AD associated differentially expressed genes, 2 genes altered in patients with AD but not in aging gene signature, and 2 inflammatory genes of interest (1 aging, and 1 non-aging gene). This systematic gene selection resulted in a total of 40 genes analyzed (Figure 1A). We also checked the association of these 40 AD related genes and known genesets (Hallmark genesets and KEGG pathways [30, 31]). These genes were found to be enriched in mainly immune response related genesets including IL-6 signaling, IFN-γ response, complement, chemokine signaling pathway, and cytokine-cytokine receptor interaction which have been suggested to be associated with AD (Figure 1B-C) [32-35]. The 40 genes were also enriched in Alzheimer’s disease in KEGG pathways as expected (Figure 1C).

### 3.2. Validation of Aging Genes

In an independent cohort from the Yale ADRC, 8 genes (20%) out of 40 were differentially expressed in peripheral blood among the CN, MCI, and dementia groups (Figure 2, see Supplementary Figures S1.1-S1.7. for all other gene expression plots). Five of the eight genes were top 9 IL-7Rα^low^ EM CD8^+^ T cell aging genes [15], including *FGFBP2, GZMH, NUAK1, PRSS23*, and *TGFBR3* (Figure 2A). The gene encoding for fibroblast growth factor binding protein 2 *(FGFBP2*) was found to be more highly expressed in the MCI group compared to the CN group (*P*-value = 0.048) and dementia group (*P*-value = 0.018). *GZMH*, a gene which encodes for the human serine protease granzyme H, was more highly expressed by the MCI group compared to the CN group (*P*-value = 0.041). *NUAK1* (gene which encodes for a protein kinase), *PRSS23* (gene that encodes for a serine protease of the trypsin family), and *TGFBR3* (gene that encodes for the transforming growth factor (TGF)-beta type III receptor) were found to have high levels of expression in the MCI group compared to the dementia group. These findings indicate increased expression of four of the top 9 IL-7Rα^low^ EM CD8^+^ T cell-associated aging genes in individuals with MCI as compared to those with dementia.

Two non-aging genes associated with AD (per both MsigDB and AD microarray datasets) [21, 22, 24] and the “Memory” dataset [23] were found to be differentially expressed (Figure 2B). *CD163* encodes scavenger receptor cysteine-rich type 1 protein M130 expressed by monocytes and macrophages. This gene was found to be differentially expressed among the three clinical groups, though adjusted *P-*values generated from post-hoc multiple comparison testing suggest no significant differences between clinical groups. *PGAP6*, which encodes post-glycosylphosphatidylinositol attachment to proteins 6, was also found to be differentially expressed with higher expression levels in the MCI group compared to both the CN (*P*-value = 0.014) and dementia (*P*-value = 0.045) groups.

Of the genes associated with the IL-7Rα^low^ EM CD8^+^ T cell aging gene signature and AD (based on the 3 publicly available AD datasets examined), *PADI4*, which encodes peptidyl arginine deiminase 4 and is related to chromatin organization and protein citrullination by converting arginine residues to citrulline residues, was found to be differentially expressed among the three clinical groups during one-way ANOVA testing though adjusted *P*-values from post-hoc multiple comparison testing suggested no significant differences in gene expression between clinical groups (Figure 2C).

Additional analysis used GLM to adjust for age, race, and sex. A full-factorial model with all possible variables, covariates, and interactions (i.e., age, sex, race, diagnosis and all interaction terms) was generated, and predictors that were not significant were removed from the predictive model. Four genes (*FGFBP2, PRSS23, TGFBR3, NUAK1*) out of 5 top IL-7Rα^low^ EM CD8^+^ T cell-associated aging genes remained differentially expressed at the *P* < 0.05 significance level (Supplementary Table S2) with decreased relative gene expression in the dementia group compared to the MCI group (Supplementary Table S3). Further analysis of the genes differentially expressed in the public AD or memory datasets identified *PGAP6*, which is up-regulated in the MCI group compared to the dementia group (adjusted *P*-value = 0.029), and *AKAP13* (encodes A-kinase anchoring protein 13) which was differentially expressed between the CN and dementia groups (adjusted *P*-value = 0.023).

### 3.3. Clinical Subgroup Clustering According to Expression of Aging Genes

To uncover potential associations between aging genes and AD status, principal component analysis (PCA) was performed using Z-scores calculated from the expression levels of all 40 genes of interest (Supplementary Figure S2). Minimal separation or clustering based on disease status was observed. Unbiased hierarchal clustering (Supplementary Figure S3) was also performed to reveal trends of gene expression in different gene target groups which demonstrated substantial portions of the MCI group and a small part of the dementia group clustered together with high expression levels of the top aging genes associated with IL-7Rα ^low^EM CD8^+^ T cells (represented by the “Top 9 IL-7Rα^low^ CD8^+^ T cell aging” gene category in Supplementary Figure S3, the cluster indicated by a blue arrow), though there was more variability and heterogeneity in clustering of the rest of the participants.

A variable loading plot was also generated (Supplementary Figure S4), mapping variables according to their contributions to PC1 and PC2. Notably, top IL-7Rα^low^ CD8^+^ T cell aging genes *SYT11, OSBPL5, PRSS23, TGFBR3, NUAK1*, and *CX3CR1* clustered together in the same quadrant and vector suggesting high correlation. The rest of the top IL-7Rα^low^ CD8^+^ T cell aging gene targets including *FGFBP2, GZMH*, and *NKG7* were less correlated but still clustered within the same region and exhibited similar contributions to PC1 and PC2 as the other T cell associated aging gene targets though with more loading on PC2.

Subgroup analyses were conducted to determine if there were patterns of clustering or separation within the clinical groups based on gene expression using PCA and unbiased hierarchal clustering. PCA analysis showed three distinct subgroups in the dementia group (Figure 3A) although no clear clustering was found among the CN and MCI groups (Supplementary Figure S2). A pattern of lower relative gene expression in the Cluster 1 dementia subgroup compared to Clusters 2 and 3 was also observed in dot plots (Figure 3B) and unbiased hierarchal clustering (Figure 3D). Notably, there was a relative increase in gene expression levels of the top IL-7Rα^low^ EM CD8^+^ T cell aging genes in Clusters 2 and 3 compared to Cluster 1 as demonstrated in Figure 3D. To evaluate cognitive functioning based on participant clustering, MoCA and CDRsob scores were plotted according to dementia subgroup clusters (Supplementary Figure S5). As Clusters 2 and 3 exhibited similar gene expression profiles (Figures 3B and 3D), both clusters were combined into one cluster which demonstrated significantly different MoCA scores from participants in Cluster 1 (*P*-value = 0.034, see Figure 3C).

## 4. DISCUSSION

AD is a disabling disease that is heterogeneous and multifactorial in its etiologies. This limits any one AD model from being globally representative of disease pathogenesis and the mechanisms that underly it [36]. This difficulty in fully characterizing AD pathology drives the ongoing exploration into several potential pathways that may contribute to the development of AD. Studies of inflammatory pathways and their association with AD have been mounting in the past decade. Evidence of systemic immune dysregulation in AD is growing, though there is still a relative paucity of studies investigating the role of adaptive immunity, including T cells, in AD [37]. Addressing the latter point is critical in that one of the most prominent changes with human aging is expansion of memory CD8^+^ T cells with senescent characteristics in peripheral blood [9]. As AD incidence rates increase with age (annual incidence of AD appreciably increase above 75 years of age [38, 39]), it is possible that senescent T cells may contribute to AD progression through altered immune activity. This study aimed to expand upon contemporary notions of adaptive immune functioning in AD by investigating the potential presence and influence of these senescent immune cells on AD through transcriptomic analysis. The results of our study show the presence of an altered CD8^+^ T cell age-associated gene signature in the peripheral blood of AD patients, warranting further studies investigating biological implications of CD8^+^ T cells, especially highly cytotoxic and inflammatory IL-7Rα^low^ EM CD8^+^ T cells, in AD.

Cross-sectional transcriptomic data from 121 age-matched ADRC participants included in this study describe alterations in several genes associated with aging and AD. Notably, 5/9 (55.6%) of the top aging genes that have been previously associated with IL□7Rα^low^ EM CD8^+^ T cells [15], a T cell population characterized by distinct senescent markers that clonally expand with age, were found to be differentially expressed in AD participants. These top aging genes include genes related to cytotoxic end products (*FGFBP2, GZMH, PRSS23*) as well pathways that have been associated with cell survival and cell cycle regulation (*PRSS23, NUAK1*) [40, 41]. After adjusting for race, sex, and age using GLM analysis, 4/9 of these top aging genes (*FGFBP2, NUAK1, PRSS23, TGFBR3*) remained differentially expressed at a significance level of 0.05. Of note, these genes have been previously associated with tumor growth and metastasis (*NUAK1, TGFBR3, FGFBP2*) [42, 43], as well as autoimmune disorders (*PRSS23*) [44]. In relation to AD, *NUAK1* overexpression is suspected to promote tau hyperphosphorylation based on findings in AD mouse models [45]. Interestingly, these genes appeared more highly expressed in the MCI group compared to the CN and dementia groups. Similar patterns have been observed in a previous study which found increases in myeloid-derived suppressor cells and FOXP3^+^ CD4^+^ regulatory T cells in the peripheral blood of MCI participants compared to age-matched healthy and mild dementia participants [46]. This pattern could be reflective of increasing immune activity and an initial anti-inflammatory response early in the disease course before the shift to full suppression of inflammation found in neurodegeneration [46].

Further, subgroup analyses of the clinical groups revealed distinct clustering and separation of dementia participants in PCA and unbiased hierarchal clustering. Dementia participants that were separated into Clusters 2 and 3 were noted to exhibit higher gene expression levels of the T cell associated aging genes compared to Cluster 1 participants. Interestingly, Clusters 2 and 3 also exhibited higher MoCA scores suggesting a higher level of cognitive functioning in participants exhibiting higher relative T cell associated aging gene expression than in participants with a more progressed dementia syndrome. This may be reflective of the heterogeneity in AD disease progression with a subset of dementia participants that could also be experiencing altered immune functioning. A recent study reported the expansion of CD45RA^+^ EM CD8^+^ T cells in the cerebrospinal fluid of patients with dementia or MCI due to AD, as well as the correlation of such cell expansion in peripheral blood with cognitive scores [16]. CD45RA^+^ EM CD8^+^ T cells expanded in the CSF were IL-7Rα^low^. These findings support the results of our study revealing the possible relationship of the IL-7Rα^low^ EM CD8^+^ T cell aging signature with cognitive function in the dementia group.

## 5. CONCLUSIONS

Altogether, the gene expression patterns in this AD cohort are suggestive of an altered immune response compared to healthy normal aging. Critically, there is significant differential gene expression of several aging genes associated with IL□7Rα^low^ EM CD8^+^ T cells, and expression patterns of such genes could divide individuals with dementia due to AD into groups with different levels of cognitive functioning. Taken together, our findings allude to the potential impact of peripheral immune dysregulation in contributing to neuroinflammatory processes associated with AD pathology, warranting further investigations of these transcriptomic findings in AD.

## 6. LIMITATIONS

There are several limitations of this study. Participants were matched according to chronological age to minimize confounding due to this variable. However, it is unlikely that this would reflect biological age; thus, affecting the potential homogeneity age-matching could produce across participants. Additionally, the heterogeneity inherent in the CN and MCI groups, as well as the low participation rates of under-represented peoples in the MCI and dementia groups, limits the generalizability of these results. Though interesting trends were observed, the dementia subgroup analysis was underpowered to reveal significant associations of the T cell associated aging gene signature with clinical correlates, such as the MoCA and CDR, among individual clusters. Thus, based on PCA and unbiased hierarchal clustering observations, Clusters 2 and 3 were combined to determine if participants with higher levels of T cell associated aging gene signatures differed from Cluster 1 participants in cognitive functioning. Lastly, it is uncertain if peripheral blood gene expression is reflective of immune alterations found in the CNS.

## ACKNOWLEDGEMENTS

We wish to thank the research participants for their contributions, and the staff of the Yale ADRC (P30-AG047270-01) and AD Research Unit for their assistance in organizing and providing the biospecimens, and associated demographics and clinical data. This research was supported by the Yale Geriatric Clinical Epidemiology and Aging Related Research T32 Training Grant (T32AG1934 to JJY) and Yale Rheumatology T32 Training Grant (5T32AR007107-46 to JPY), National Institute on Aging (P30AG066508 to AM, HA, CvD), Claude D. Pepper Older Americans Independence Center at Yale School of Medicine, funded by the National Institute on Aging (P30AG021342 to HA) and the National Institutes of Health (1R01AG056728 and R01AG055362 to IK). The funders had no role in the design and conduct of the study; collection, management, analysis, and interpretation of the data; preparation, review, or approval of the manuscript; and decision to submit the manuscript for publication.

## DISCLOSURES

Parts of this work have been submitted in partial fulfillment of the requirements for the Master of Health Science degree to be conditionally awarded by the Yale School of Medicine to JJY.

## CONFLICTS OF INTEREST

Authors declare no conflict of interest.

## AUTHOR CONTRIBUTIONS

JJY, HJP, and JPY designed the study, performed the experiments, analyzed and interpreted the results, and participated in writing the manuscript; JJY, HHB, CvD, and APM participated in clinical consensus diagnosis meetings that involved diagnosis of ADRC participants as well as characterizing and organizing of clinical data associated with biospecimens utilized in this research study; MK and SY retrieved, analyzed, and organized publicly available gene expression datasets utilized to identify gene targets of interests investigated in this study; HSK, MSS, SU, and HA analyzed and interpreted the results, as well as participated in writing the manuscript; HA, APM, CvD, RB, and SY interpreted the results and participated in writing the manuscript; IK designed the study, analyzed and interpreted the results, participated in writing the manuscript and supervised the research.

## Notes

### Competing Interest Statement

The authors have declared no competing interest.

