## Supplementary material for "“Altered CD8^+^ T cell associated aging gene signature in the peripheral blood of patients with Alzheimer’s disease”": Tables

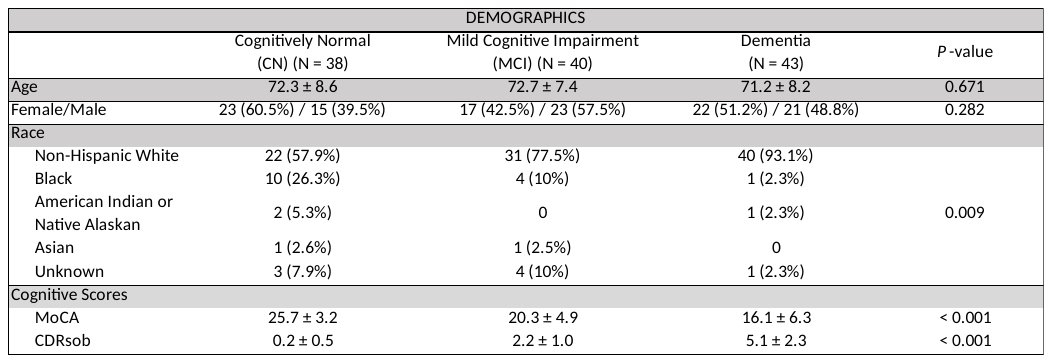

**Table 1. Demographics of Study Participants**

Demographic characteristics of 121 Alzheimer’s Disease Research Center (ADRC) participants involved in this study and organized according to clinical group. There was a predisposition of MCI and dementia cohorts to contain an unequal distribution of participants based on race (χ^2^(8) = 20.2; *P*-value = 0.009). Abbreviations: MoCA, Montreal Cognitive Assessment; CDRsob, Clinical Dementia Rating sum of boxes
