## Supplementary material for "“Altered CD8^+^ T cell associated aging gene signature in the peripheral blood of patients with Alzheimer’s disease”": Figures

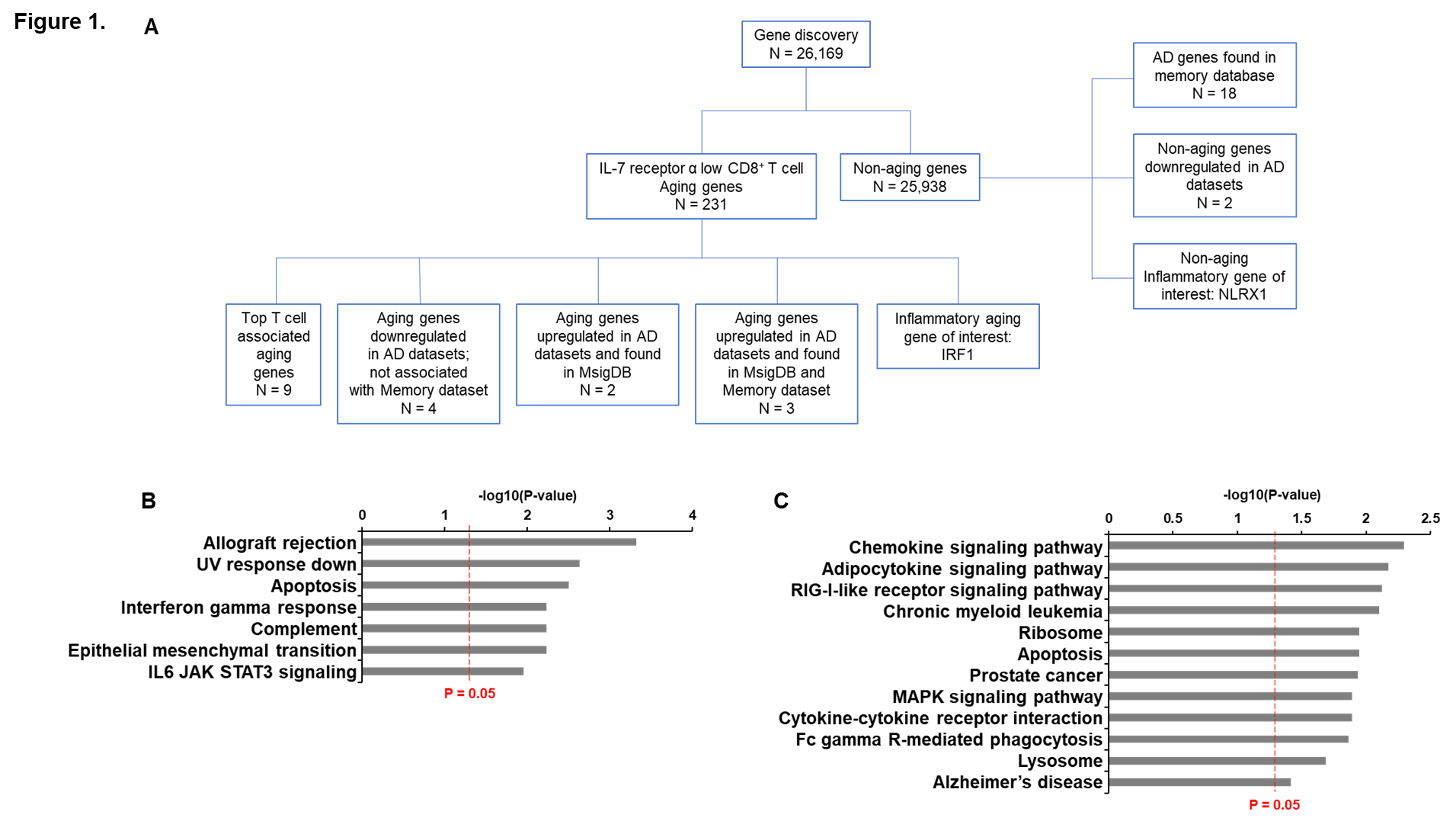

**Figure 1. Gene discovery diagram and associated Hallmark genesets and KEGG pathways**

(A) Gene discovery involved the analysis of 26,169 total genes based on publicly available databases that included AD datasets (GSE140829, and GSE63060 and GSE63061) and memory transcriptomic data (GSE127711) as well as the Molecular signatures database (MSigDB). This resulted in the analysis of 19/231 aging genes as well as 21 genes of interest associated with AD and/or inflammation. (B-C) Hallmark (B) and KEGG (C) genesets pathways enriched with 40 AD associated genes.

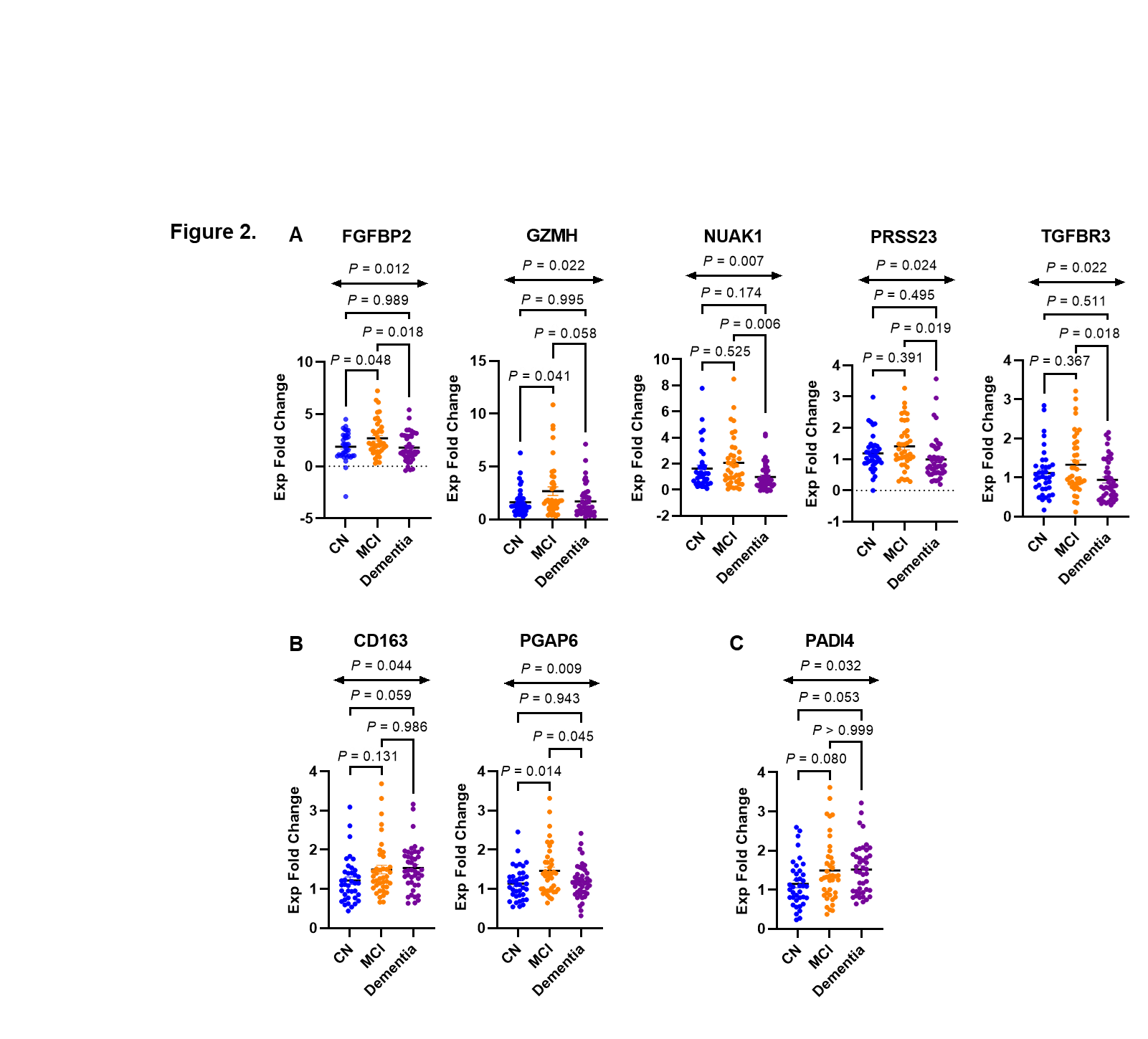

**Figure 2. A set of genes associated with IL-7 receptor alpha low effector memory CD8^+^ T cells and Alzheimer’s disease (AD) is differentially expressed in peripheral blood of patients with AD.**

Reverse transcription qPCR (RT-qPCR) analysis showing differentially expressed genes in peripheral blood of cognitively normal (CN) and AD participants with mild cognitive impairment (MCI) or dementia. (A) top IL-7 receptor alpha low (IL-7Rα^low^) effector memory (EM) CD8^+^ T cell-associated aging genes. (B) genes associated with memory or AD datasets but not in the T cell associated aging gene signature. (C) *PADI4,* an aging gene found to be upregulated in 3 AD datasets. *P* values were obtained by ANOVA and adjusted during post-hoc multiple comparison testing.

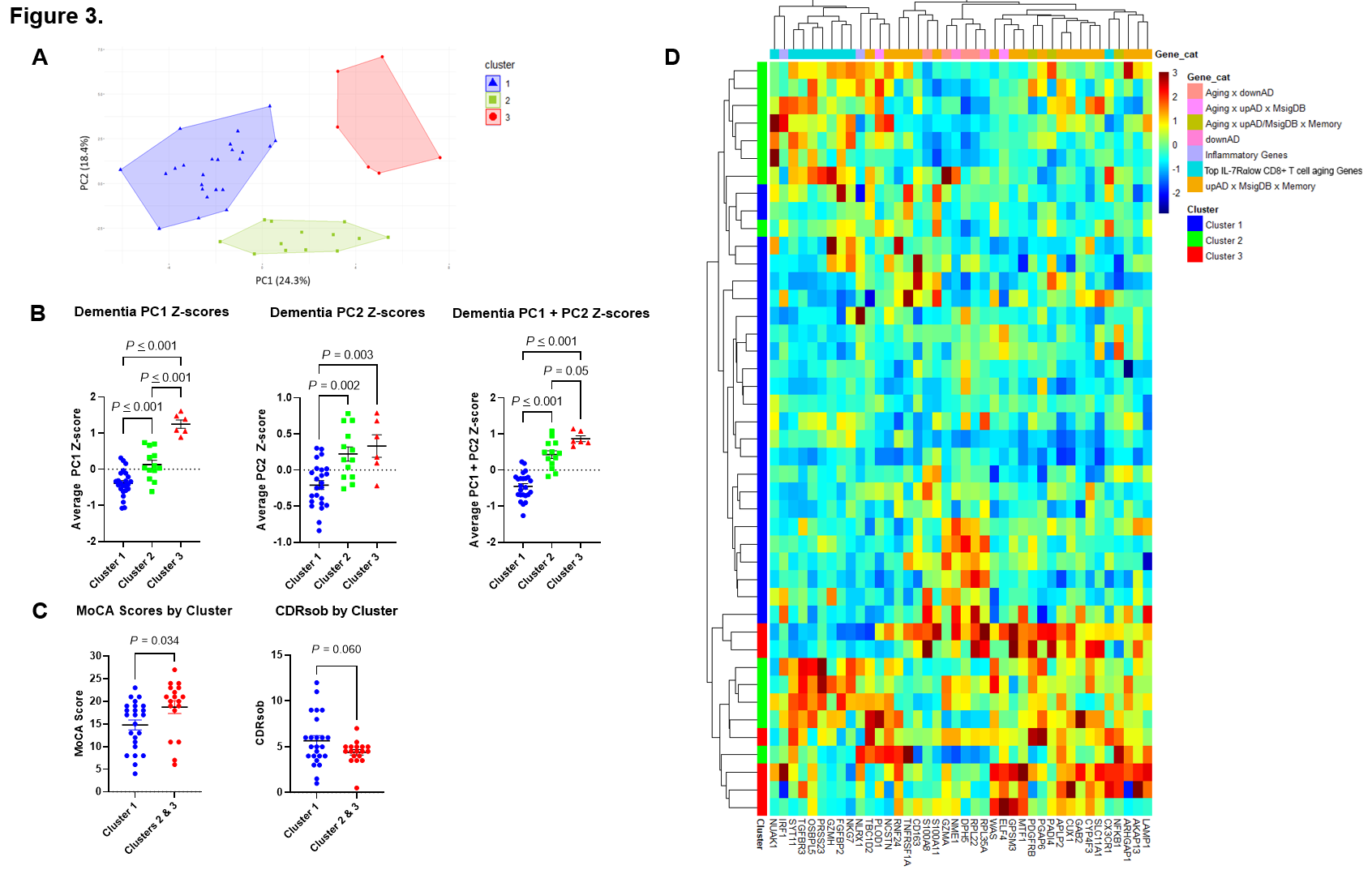

**Figure 3. Unbiased PCA and hierarchal clustering of dementia subgroups demonstrate differential expression of IL-7 receptor alpha low effector memory CD8^+^ T cell associated aging genes based on measures of cognitive functioning.**

Principal component analysis (PCA) based on the expression levels of genes in peripheral blood of AD participants was done. (A) PCA yielded three clusters: Cluster 1 (blue triangles), Cluster 2 (green boxes), and Cluster 3 (red circles). (B) Average Z-scores of dementia subgroup clusters based on principal component (PC) loading. (C) MoCA and CDRsob scores in Cluster 1 vs. Clusters 2 & 3. (D) Unbiased hierarchal clustering gene expression heatmap of dementia subgroups categorized impartially according to cluster designation and gene target category; MoCA, Montreal Cognitive Assessment; CDRsob, Clinical Dementia Rating scale sum of boxes. *P* values were obtained by ANOVA with post-hoc multiple comparison testing or Welch’s *t*-test.
