## Supplementary Materials for "“Altered CD8^+^ T cell associated aging gene signature in the peripheral blood of patients with Alzheimer’s disease”"

1. **Methods**
2. **Tables**
3. **Figures**

**1. Methods**

1.1. **RNA Isolation and Complementary DNA synthesis Protocol**

RNA was isolated using a modified QIAGEN RNeasy Kit protocol. Frozen whole blood was allowed to thaw in QIAzol Lysis Reagent (QIAGEN) and homogenized by pipetting. 1 mL of homogenate was transferred into a 1-2mL microcentrifuge tube where 200 µl of chloroform was added and the mixture shaken vigorously for 15 seconds. Afterwards, the mixture was incubated at room temperature for 2-3 minutes and then centrifuged at 10,000 RPMs (Eppendorf 5417R Centrifuge) for 15 minutes at 4˚C. After centrifugation, 400-500 µl of the upper aqueous phase of the mixture was transferred to a new microcentrifuge tube taking care to avoid the interphase. 70% ethanol was added at a 1:1 volume ratio and the mixture was vortexed. 700 µl of the resulting mixture was then transferred to a RNeasy Mini spin column (QIAGEN) and centrifuged at room temperature for 15 s at ≥8000 x g. The flow-through was discarded and repeated passing of the ethanol-sample mixture through the RNeasy Mini spin column was done until none of the original mixture was left. The DNase Digest protocol (QIAGEN) was then conducted to purify the sample RNA. Afterwards, the sample RNA was washed with 500 µl Buffer RPE two times and dried by centrifuging at full speed (≥ 13,000 RPM) for 3-5 min. The RNeasy Mini spin column was placed in a new microcentrifuge tube and 30 µl of RNase-free water was added and left to rest for 1 minute before centrifuging for 1 min at ≥ 10,000 RPM to collect the resulting RNA solution. The RNA solution was placed in an ice bath and was subsequently tested for quality and quantity via the NanoDrop™ 2000 Spectrophotometer. RNA templates of good quality and quantity were then utilized for complimentary DNA (cDNA) synthesis utilizing the iScript cDNA synthesis kit protocol (Bio-Rad).

1.2. **Gene Discovery**

Forty genes of interest were identified from publicly available databases based on their associations with at least one transcriptomic dataset or their relation to significant inflammatory or aging markers. Datasets that were utilized in gene discovery include microarray expression profiles (GSE140829.GPL15988 [1] and GSE63063.GPL6947 and GLPL10558 [2]) derived from total RNA obtained from the peripheral blood of dementia patients. Additional genes were obtained from the molecular signature database (MSigDB) [3] which included over 6,700 gene sets originating from a diverse set of sources and tissue types. Specifically, data were obtained from tissue and microarray databases associated with Alzheimer’s disease (AD) pathways [4], as well as direct brain tissue and endothelial cell microarray data assorted by Blalock et al. [5] and Wu et al. [6]. Lastly, a blood-based biomarker database geared towards early detection of AD risk was utilized to discover genes of interest that were associated with short memory dysfunction as measured by the Hopkins Verbal Learning Test [7]. After discovery, these genes were separated into seven categories:

i. Top 9 aging genes with the highest correlation to the IL-7Rα^low^ EM CD8^+^ T cell subset previously discovered in a prior study [8].
ii. 2 aging genes that are known to be associated with AD according to the MSigDB and are upregulated in at least 2 of 3 publicly available AD datasets (GSE140829 [1] and GSE63060 and GSE63061 [2]).
iii. 3 aging genes that had false discovery rate (FDR) < 0.05 in the short-term memory study (GSE127711) [7] and known to be associated with AD according to the MSigDB and/or are altered in the at least 2 of 3 publicly available AD datasets (GSE140829, GSE63060, and GSE63061).
iv. 4 aging genes downregulated in publicly available AD datasets (GSE140829, GSE63060, and GSE63061).
v. 18 genes altered in the short-term memory study and known to be associated with AD (i.e., altered in AD according to the MsigDB and upregulated in publicly available AD datasets, GSE140829, GSE63060, and GSE63061) but are not in the aging gene signature.
vi. Non aging genes that are downregulated in any AD datasets (NME1 and RPL35A).
vii. Inflammatory genes of interest including IRF1 and NLRX1.
2. **Tables**

**Table S1. Gene primer sequences arranged by gene categories**

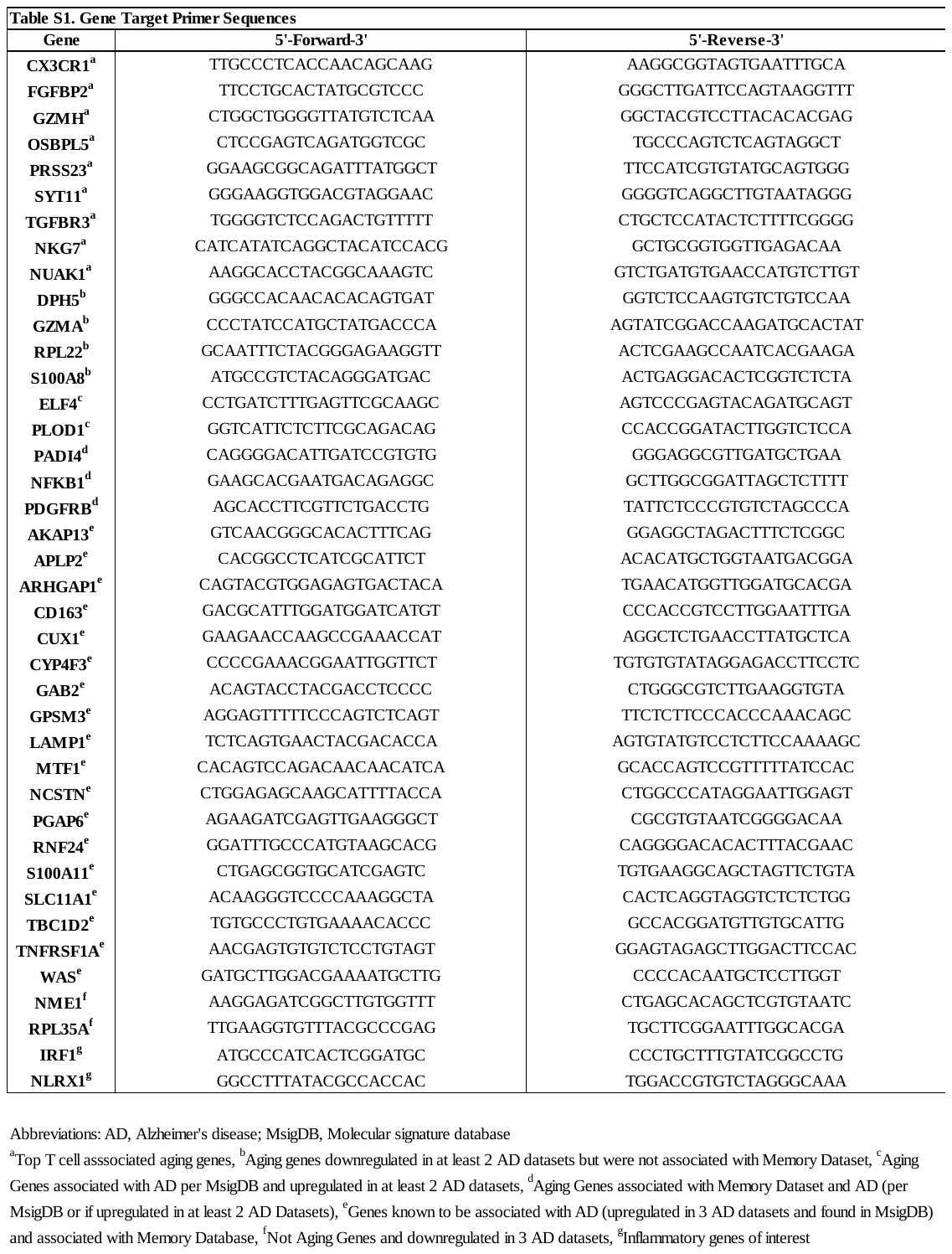

**Table S2. GLM analysis for differentially expressed genes**

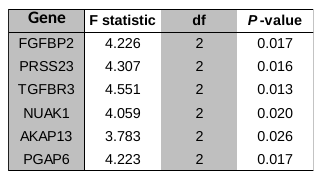

Table demonstrating F statistics and corresponding *P*-values calculated from GLMs for differentially expressed genes. GLMs were adjusted for age, sex and race, and standard error (95% CI) by clinical group. Four out of the nine top T cell associated aging gene targets (*FGFBP2*, *PRSS23*, *TGFBR3*, *NUAK1*) that were analyzed by qPCR remained differentially expressed at a significance level of 0.05.

Abbreviations: GLM, general linear model

**Table S3. LSM differences between clinical groups generated from GLM analysis for differentially expressed genes**

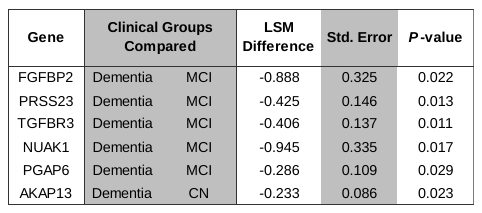

Table displaying post-hoc multiple comparison testing adjusted *P-*values after Šidák correction for differentially expressed genes suggesting lower levels of expression in the dementia group compared to the MCI group for most aging genes after adjustment (lower LSM estimations reflective of lower predicted relative gene expression). Abbreviations: GLM, General linear model; LSM, least-squares means; CN, cognitively normal; MCI, mild cognitive impairment

4. **Figures**

**Figure S1.1. Top T cell associated aging genes**

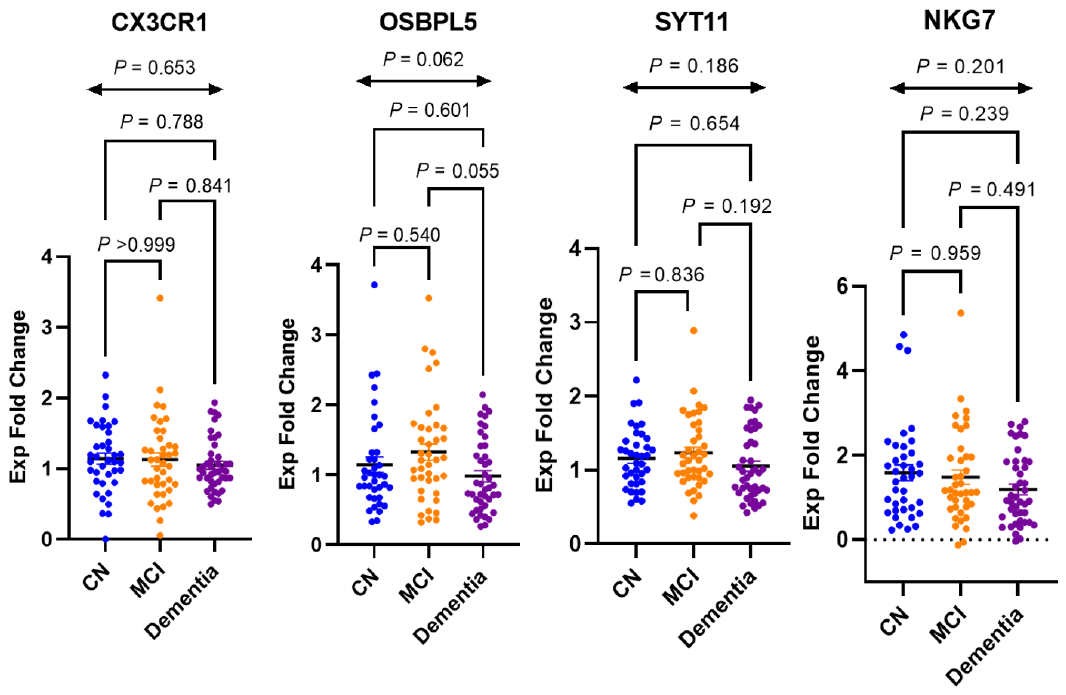

**Figure S1.2. Aging genes downregulated in at least 2 AD datasets and were not associated with Memory Dataset**

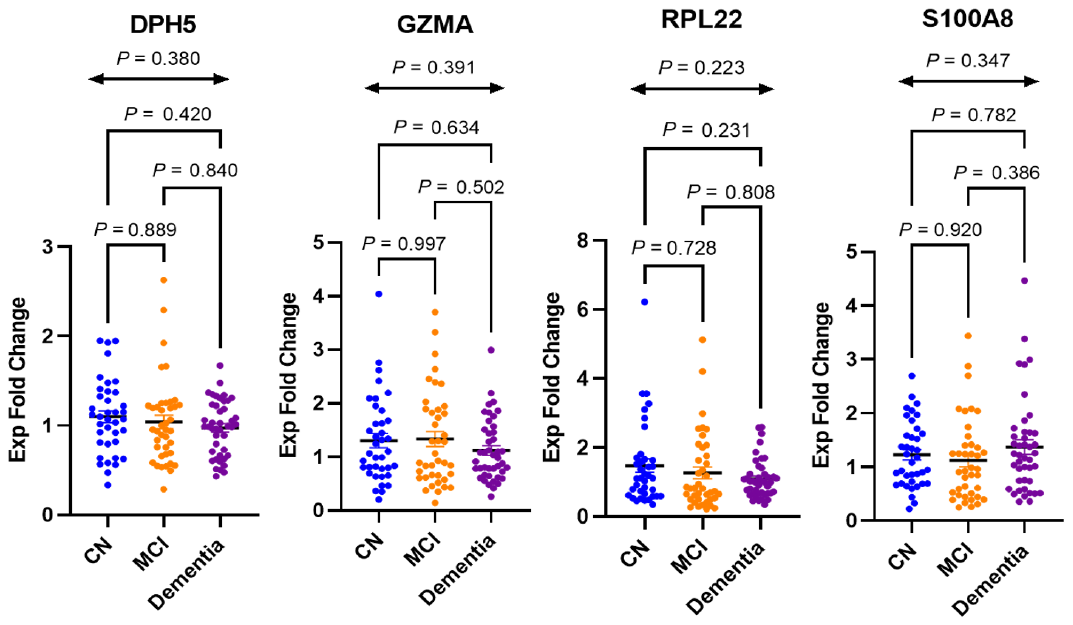

**Figure S1.3. Aging Genes associated with AD per MsigDB and upregulated in at least 2 AD datasets**

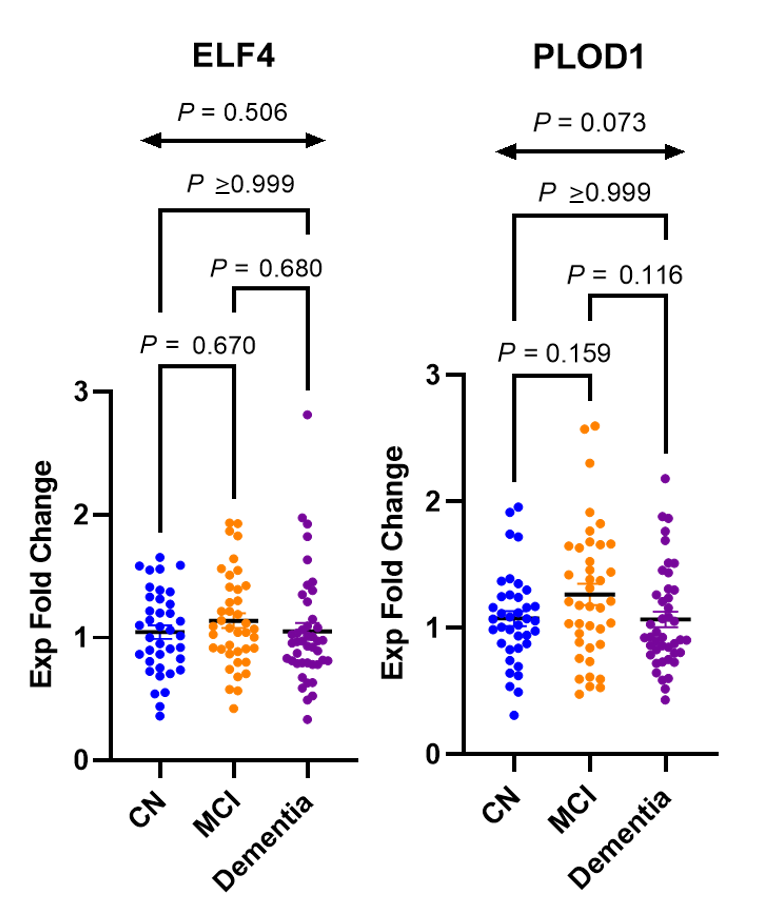

**Figure S1.4. Aging genes associated with Memory Dataset and AD per MsigDB and/or upregulated in at least 2 AD Datasets**
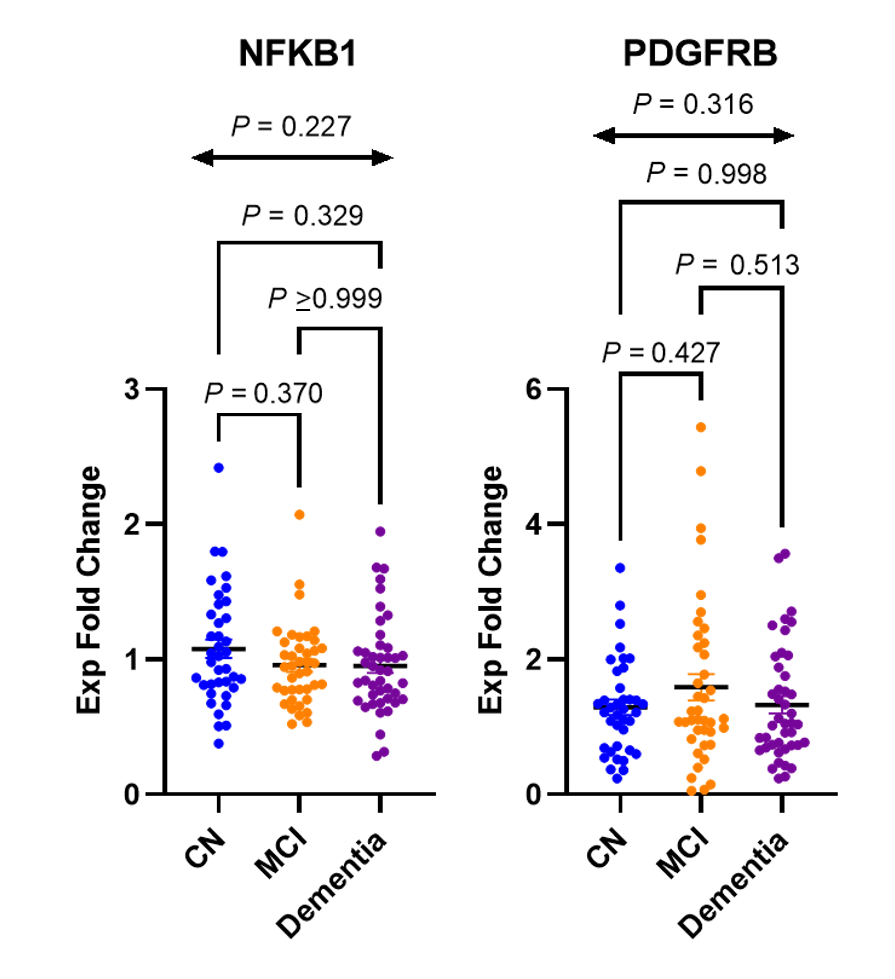

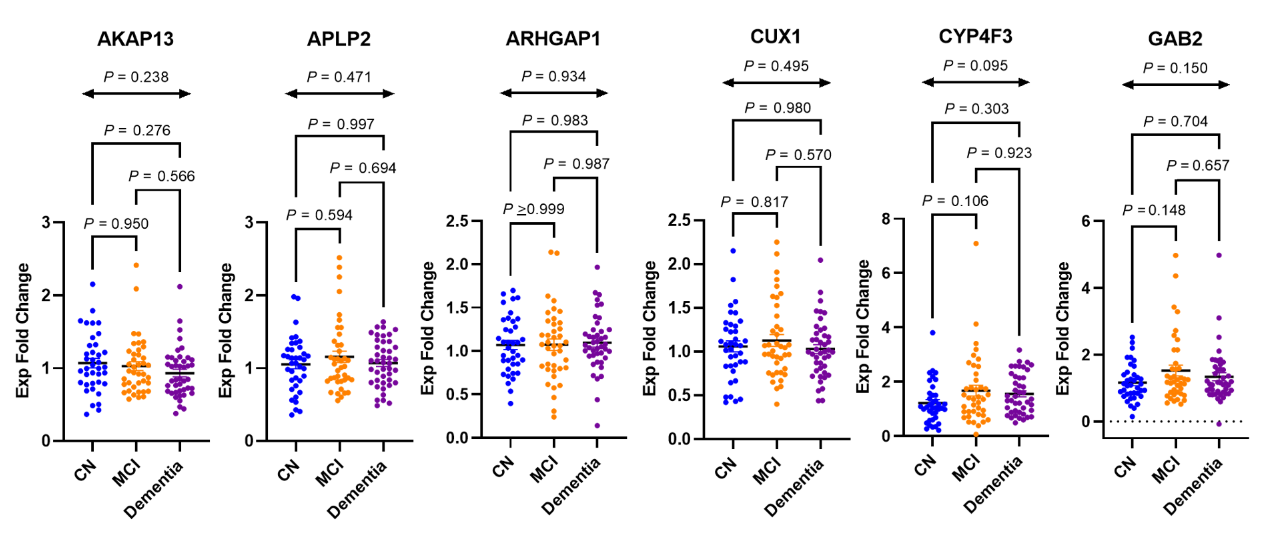
**Figure S1.5. AD genes associated with memory database**

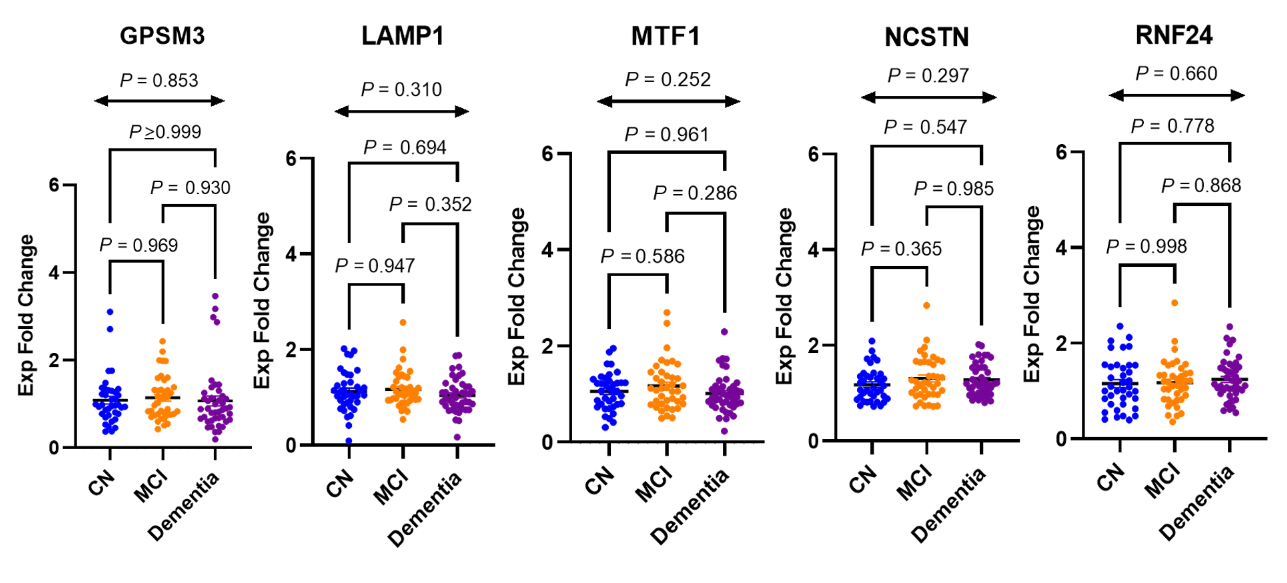
**Figure S1.5. AD genes associated with memory database (cont’d)**

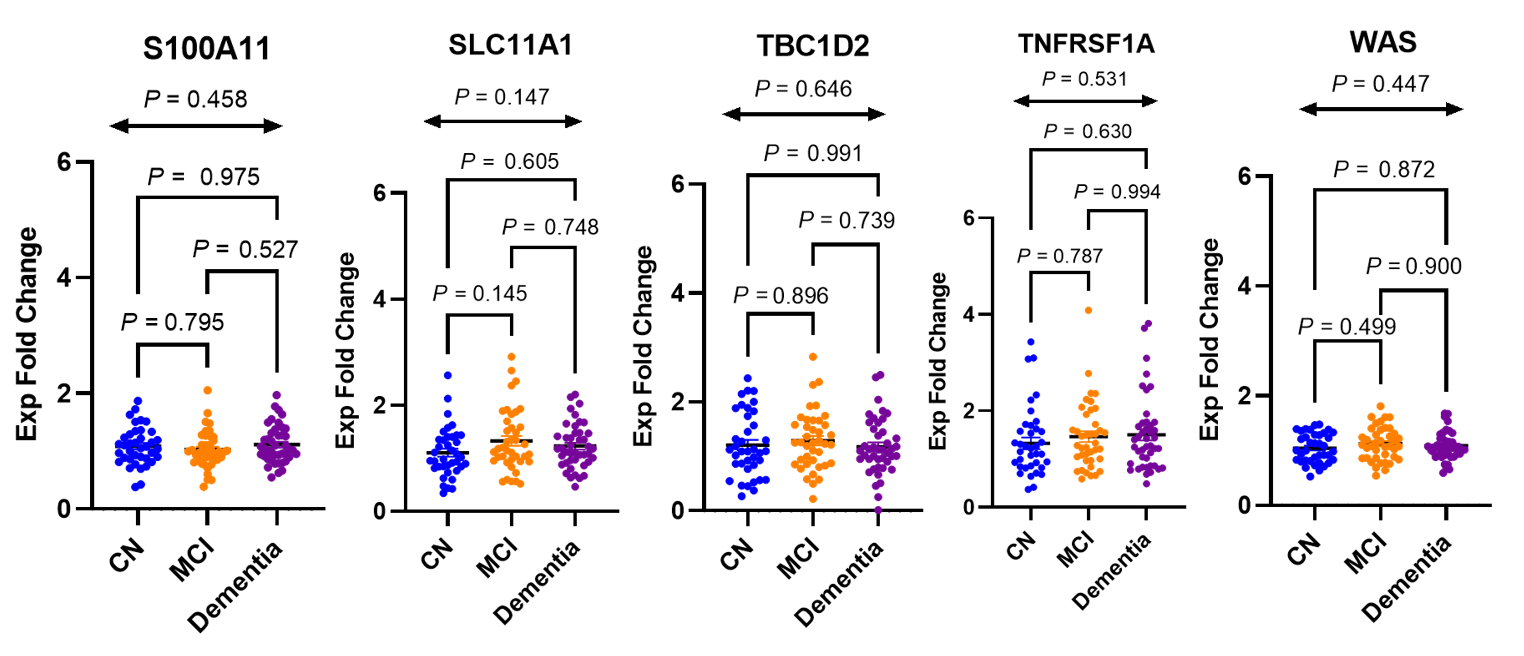
**Figure S1.5. AD genes associated with memory database (cont’d)**

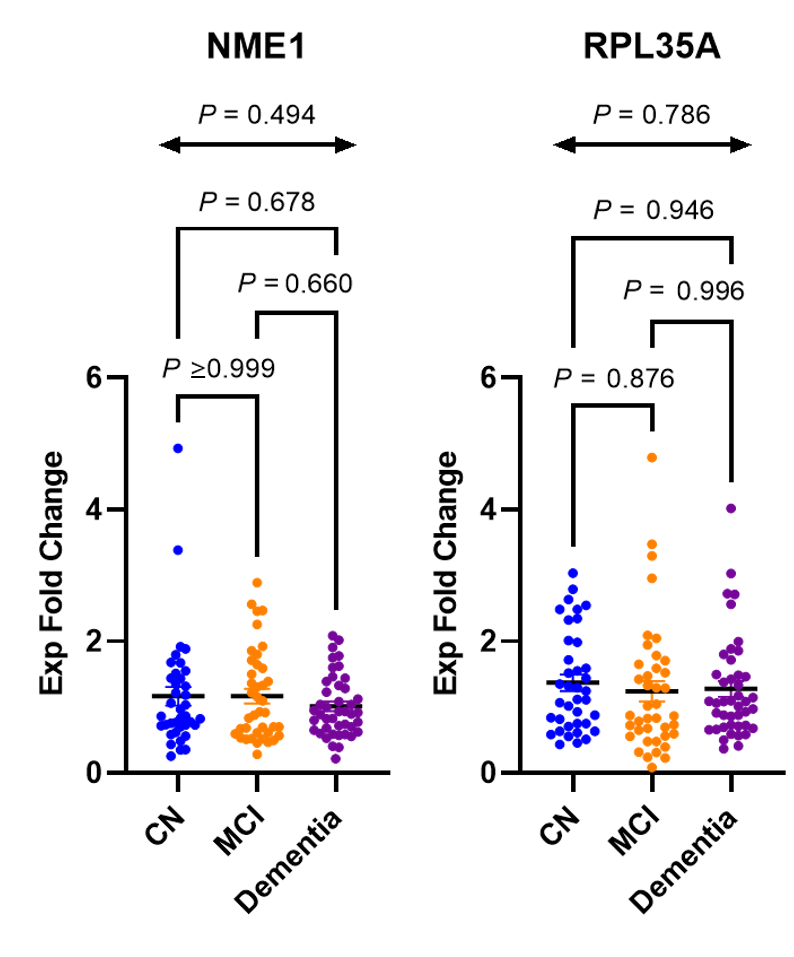
**Figure S1.6. Non-Aging Genes and downregulated in 3 AD datasets**

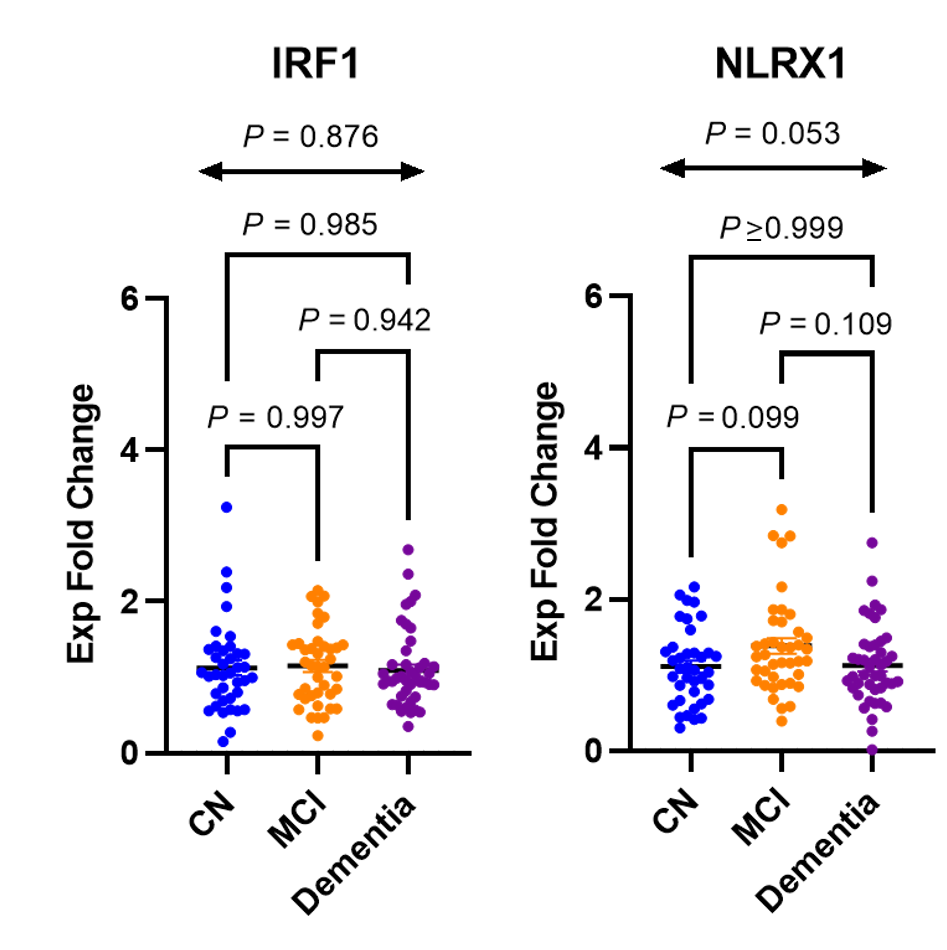
**Figure S1.7. Inflammatory genes of interest**

Figures S1.1.-S1.7. Reverse transcription qPCR (RT-qPCR) analysis showing differentially expressed genes in peripheral blood of cognitively normal (CN) and AD patients with mild cognitive impairment (MCI) or dementia. Plots were shown with one-way ANOVA *P*-values (bidirectional arrow) and post-hoc multiple comparison testing adjusted *P-*values for genes that were not found to be differentially expressed between the three clinical groups. Abbreviations: AD, Alzheimer’s disease

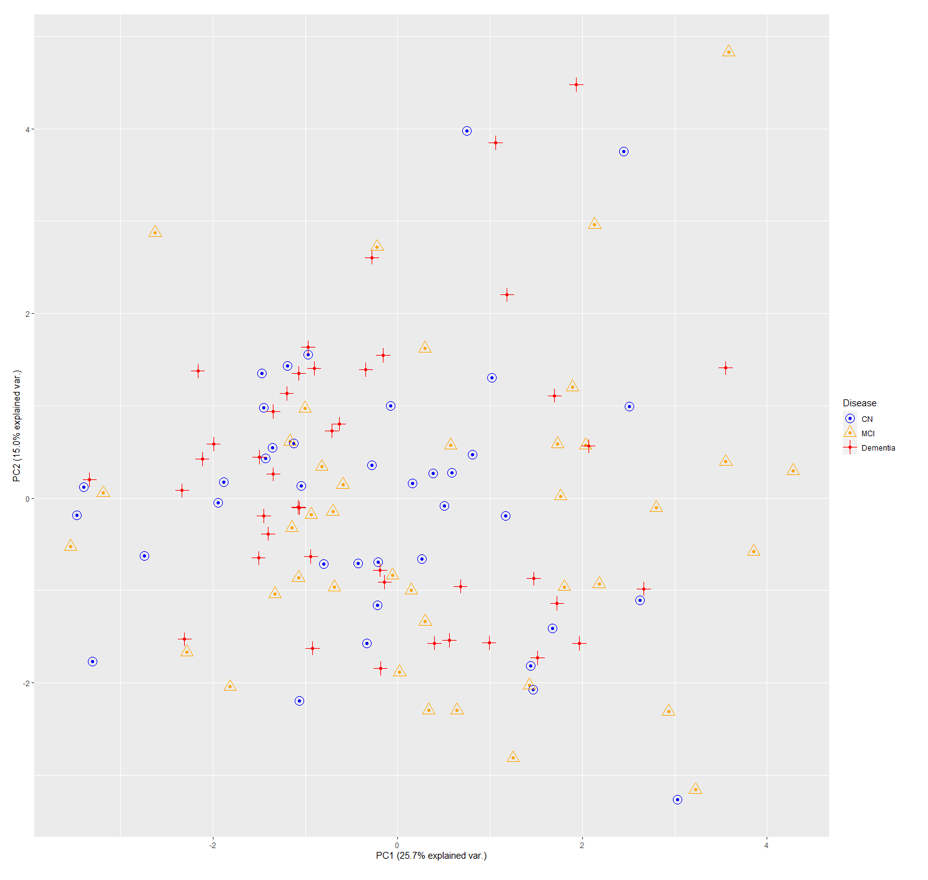

PC2 (15% explained var.)

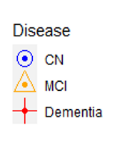

PC1 (25.7% explained var.)

**Figure S2. All Genes PCA Plot**

PCA plot generated using all gene target Z-scores calculated for all participants. Points on plot represent individual participants identified by clinical group. Blue circles represent CN, yellow triangles represent MCI, and red crosses represent dementia group participants. Abbreviations: PCA, principal component analysis; var, variance; CN, cognitively normal; MCI, mild cognitive impairment

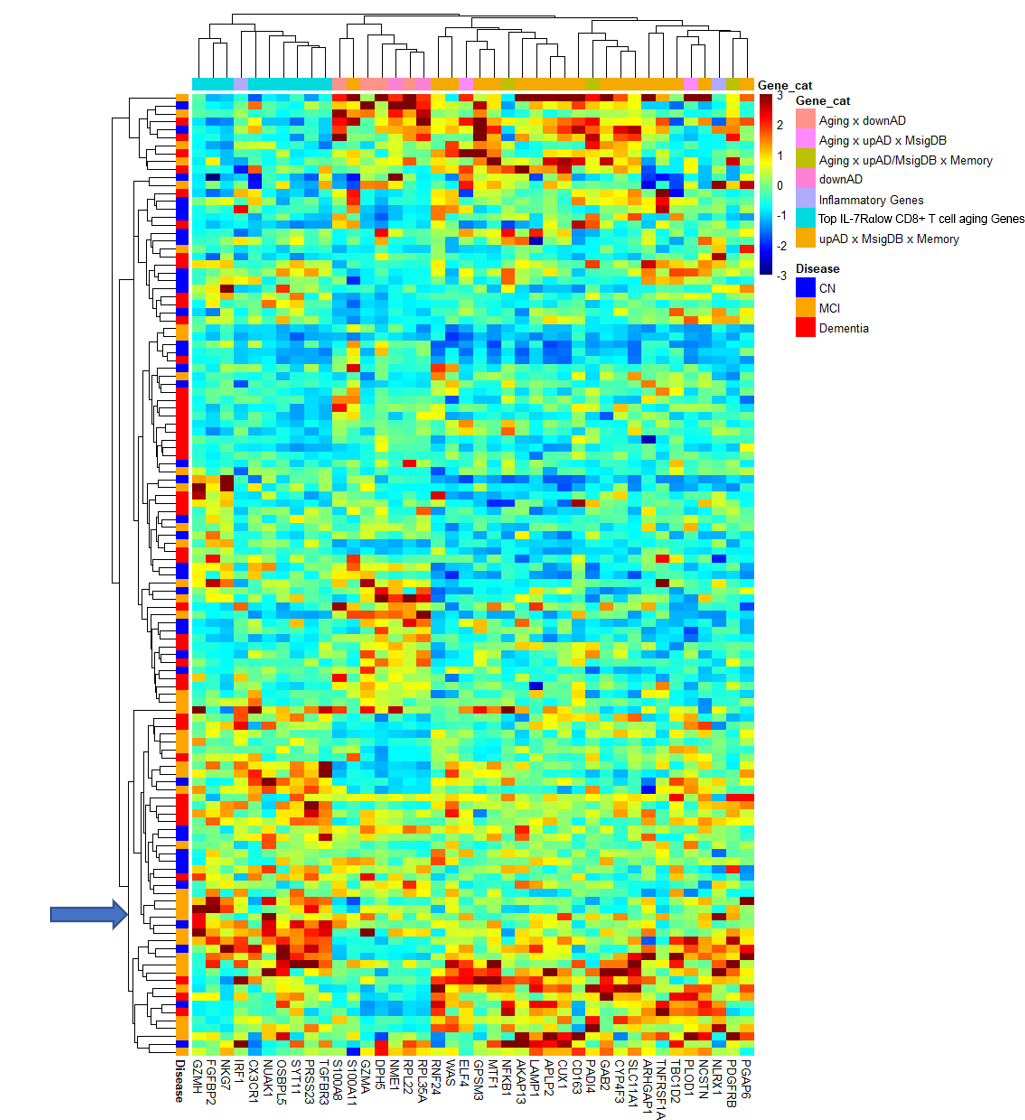

**Figure S3. Hierarchal Clustering Heatmap**

Unbiased hierarchal clustering of all gene categories according to differential gene expression of individual participants. Clustering is labeled according to gene target category on the horizontal axis and clinical group on the vertical axis. Clustering is based on the “Euclidean” distance of gene target expression levels from one another. Blue arrow indicates a cluster with high expression levels of the top aging genes associated with IL-7Rα^low^ EM CD8^+^ T cells.

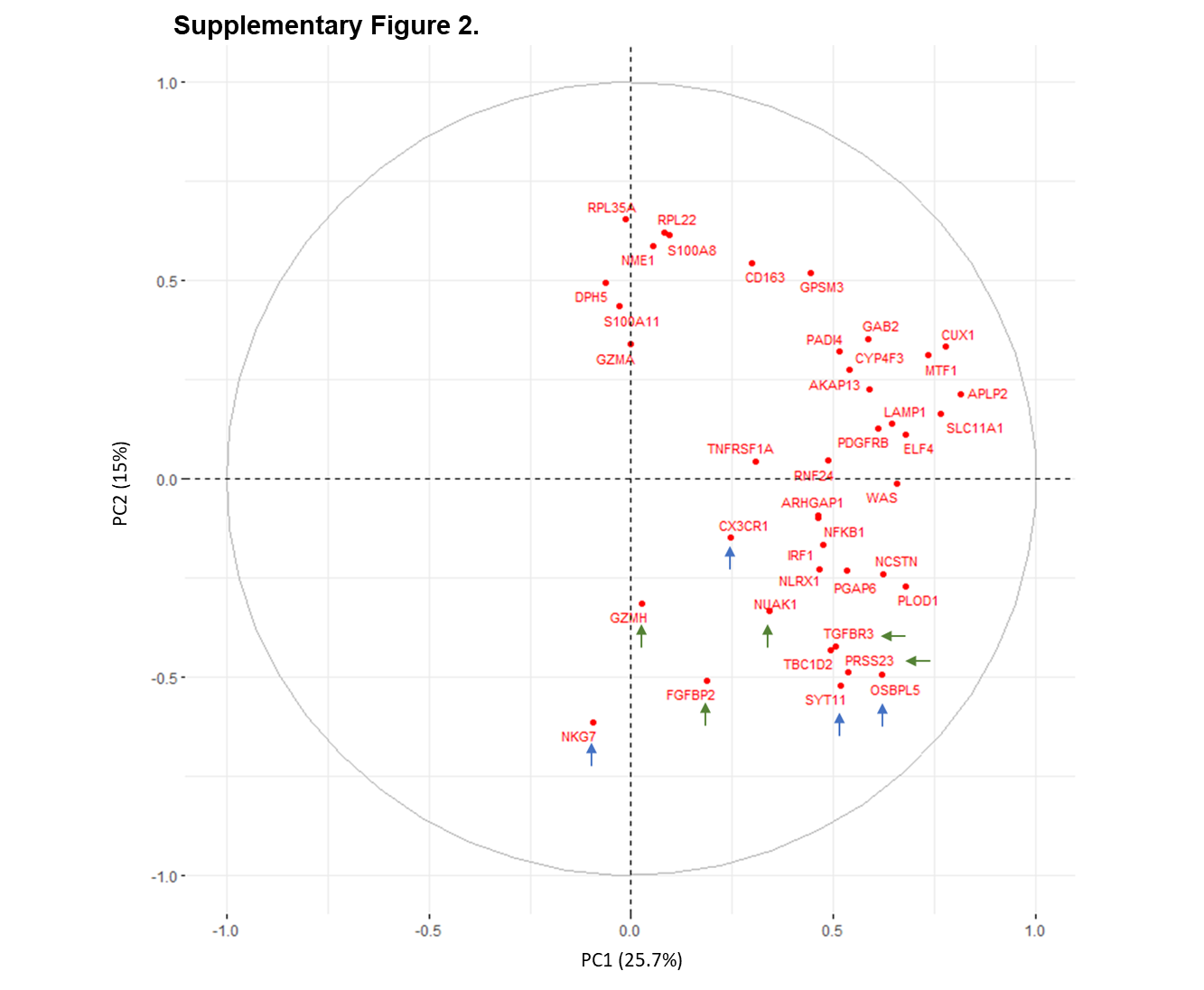

**Figure S4. All Genes Variable Loading Plot**

Loading plot visualizing gene target loading to PC1 and PC2. Green arrows point to differentially expressed top T cell associated aging genes (*P*-value ≤ 0.05). Blue arrows represent top T cell associated aging genes that were not found to be differentially expressed at a 0.05 significance level. Despite differences in significance of gene differential expression across the clinical groups, all top T cell associated aging genes appeared to cluster in the same quadrant and vector suggesting they highly correlated with one another. Abbreviations: PC, principal component

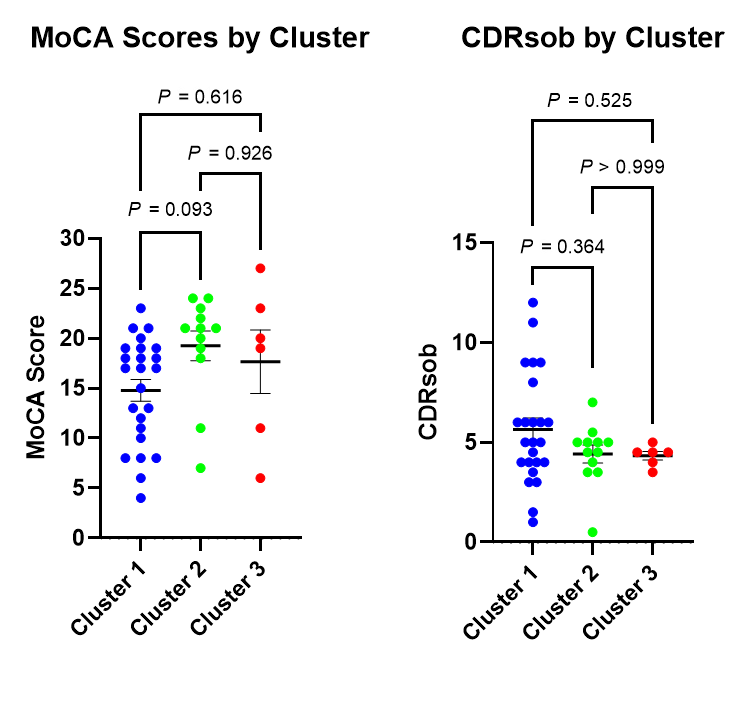

**Figure S5. MoCA and CDRsob Scores per Dementia Subgroup Cluster**

MoCA and CDRsob scores plotted according to cluster designation in the dementia group. Abbreviations: MoCA, Montreal Cognitive Assessment; CDRsob, Clinical Dementia Rating scale sum of boxes
